# Loss of DNA mismatch repair genes leads to acquisition of antibiotic resistance independent of secondary mutations

**DOI:** 10.1101/2025.06.20.660601

**Authors:** David E. Bautista, Cassidy R. Whitehead, Angela M. Mitchell

**Affiliations:** Department of Biology, Texas A&M University, College Station, Texas, USA

**Keywords:** Antibiotic Resistance, Lysis, Mismatch Repair, Stress Response, Homologous Recombination

## Abstract

Antibiotic resistant bacteria have been a rising clinical concern for decades. Beyond acquisition of alleles conferring resistance, bacteria under stress (e.g., from changing environmental conditions or mutations) can have higher intrinsic resistance to antibiotics than unstressed cells. This concern is expanded for gram-negative bacteria which have a protective outer membrane serving as an additional barrier against harmful molecules such as antibiotics. Here, we report a pathway which increases antibiotic resistance (i.e., minimum inhibitory concentration) in response to inactivation of the DNA Mismatch Repair pathway (MMR). This pathway led to increased intrinsic resistance and was independent of secondary mutations. Specifically, deletion of the DNA mismatch repair genes *mutL* or *mutS* caused resistance to various antibiotics spanning different classes, molecular sizes, and mechanisms of action in several different *E. coli* K-12 MG1655 strains, and in *Salmonella enterica* serovar Typhimurium LT2. This pathway was independent of the SOS response (severe DNA damage response). However, the patterns of resistance correlated with previously reported increases in MMR mutants in rates of homoeologous recombination, homologous recombination between non-identical DNA strands. Mutations expected to lower rates of recombination in MMR mutants also decreased the resistance to most antibiotics. Finally, we found lysis occurs in MMR mutants and may contribute to resistance to other antibiotics. Our results have demonstrated a novel mechanism that increases antibiotic resistance in direct response to loss of MMR genes, and we propose this resistance involves increased rates of homoeologous recombination and cell lysis. The increased antibiotic resistance of MMR mutants provides a path for these cells to survive in antibiotics long enough to develop more specific resistance mutations and so may contribute to the development of new clinical resistance alleles.

**Author Summary:** Antibiotic resistance has become a worldwide clinical threat and understanding resistance mechanisms is essential for continued treatment of bacterial infections. Here, we investigate a novel mechanism acting in *E. coli* and *Salmonella* when DNA mismatch repair (MMR) is lost that increases the concentration of many antibiotics needed to kill cells and does not require secondary mutations. Our data suggest increased rates of homoeologous recombination (between non-identical DNA strands) and the lysis of some cells within a population are involved in this resistance. This pathway provides a mechanism through which cells with an increased mutation rate due to loss of MMR could survive long enough in the presence of antibiotics to develop new resistance mutations, leading to the spread of antibiotic resistance.

## Introduction

Antibiotic resistance has become a global threat. Every year millions of infections occur from antibiotic resistant microbes, many of those being infections with gram-negative bacteria (1, 2). In the United States alone, the Centers for Disease Control and Prevention (CDC) estimate there are approximately three million antibiotic resistant infections, resulting in more than 36,000 deaths each year (3). With the slow development of new therapeutics and microbes constantly adapting to antimicrobial compounds, understanding how bacteria acquire resistance to antibiotics is a pressing issue.

To be defined as resistant to an antibiotic, the minimum inhibitory concentration (MIC) of the antibiotic for the strain must be above a well-defined clinical breakpoint (4). There are multiple mechanisms microbes use to become resistant to harmful substrates, including limiting drug uptake, modifying a drug target, inactivating a drug, active drug efflux, and, for some antibiotics, becoming auxotrophic (5). These mechanisms can be mediated by plasmid acquisition, chromosomal mutations, addition of new genes, or phenotypic adaptation (6). Furthermore, newly developed resistance alleles can quickly expand in populations through vertical and horizontal gene transfer, becoming widespread. On top of these genetic alterations, the outer membrane (OM) increases the intrinsic resistance of gram-negative bacteria to antibiotics (2, 7, 8). Often, mutations that decrease OM permeability can increase resistance to an antibiotic (i.e., the MIC) without crossing the clinical breakpoint to become resistant (7, 9–12). However, these mutations increase the probability of the strain acquiring alleles to become resistant (7, 9–12). In this study, we will describe a phenotype of increased resistance that occurs in DNA mismatch repair (MMR) mutants.

Nucleotide mismatches, often a result of DNA replication, cause alterations to the local structure of the DNA. MMR corrects nucleotide mismatches as well as other types of DNA damage that cause disruption of DNA structure (13–16). MMR requires the protein components MutS, MutL, MutH, DNA helicase II (aka MutU/UvrD), exonucleases, single-stranded binding protein, DNA polymerase III, DNA Ligase, and Dam methylase (17) (**Figure S1**). MutS, known as the mismatch recognition protein, recognizes base-to-base mismatches and small nucleotide insertion/deletion mispairings and is ATP-dependent. MutS then recruits MutL to form a complex. After the complex is formed, MutH is recruited and scans 5′ or 3′ to a hemi methylated GATC site. Then, it makes a strand-specific nick of the newly synthesized DNA strand (18). DNA helicase (UvrD) then unwinds the strand with the nick, followed by degradation of the strand by an exonuclease. Degradation begins at the nick and continues past the mismatch that was recognized by MutS. The single strand will then be resynthesized and ligated by DNA polymerase III and DNA ligase, respectively. Finally, the new strand will be methylated, completing the repair process (15).

Point mutations to MMR genes have been reported in antibiotic resistant clinical isolates of both gram-positive and gram-negative bacteria (19–25). These strains also had other mutations that were thought to be the cause of the antibiotic-resistant phenotype, with MMR mutations facilitating development of the other mutations due to the lack of strand specificity for mismatch repair by other DNA repair pathways. However, the effect of the MMR mutations on antibiotic resistance in isogenic strains was not evaluated.

In a study characterizing an OM permeability mutant (Δ*yhdP*), Mitchell, *et al.* observed that disruption of MMR genes by transposon insertion could cause vancomycin resistance in *E. coli* K-12 (26). YhdP is a protein which plays a role in phospholipid transport between the inner membrane (IM) and OM (27–29). Deletion of *yhdP* causes cells to become much less resistant to vancomycin and SDS EDTA (26). Transposon mutagenesis was performed in a strain with a deletion of *yhdP.* Transposon mutants with increased resistance to vancomycin were selected from the transposon library. Transposon insertions in *mutL* or *mutS*, inactivating MMR, led to vancomycin resistance in the Δ*yhdP* strain; however, no effect of these mutations was observed for Δ*yhdP*’s SDS EDTA sensitivity, suggesting that inactivating MMR affected vancomycin resistance rather than Δ*yhdP* specific functions (26).

Here, we set out to explore the link between inhibition of MMR and increased vancomycin resistance in *E. coli* K-12. We have found the resistance extends to many antibiotics beyond vancomycin and is caused directly by loss of MMR without the need for further mutations. It can be observed in wild-type *E. coli*, in OM permeability mutants, and in *Salmonella enterica* Serovar Typhimurium. Furthermore, our data suggest the mechanism of this resistance involves increased rates of homoeologous recombination (recombination between non-identical DNA strands) and increased rates of cell lysis. Understanding mechanisms of increased resistance via inhibition of MMR will allow for new discoveries to be made on how to target antibiotic resistance mutants and alleviate the epidemic of antibiotic-resistant infections.

## Results

### Increased antibiotic resistance occurs directly from loss of MMR genes and not from increased mutation rates

Hypermutators, such as cells with inhibition of MMR function, have been linked to antibiotic resistance (30). Clinical antibiotic-resistant strains with MMR mutations have been assumed to have increased resistance rates allowing them to gather other favorable mutations, but the direct effects of MMR on antibiotic resistance have not yet been studied. Given that we previously observed increased vancomycin resistance in a more vancomycin sensitive Δ*yhdP* strain with several independent transposon mutants in MMR genes in *E. coli* K-12 (26), we first investigated whether the relationship between loss of MMR and increased vancomycin resistance required secondary mutations facilitated by increased mutation rates. If the increased antibiotic resistance was due to secondary mutations, the level of resistance should differ between separately constructed MMR mutants as these strains would have different spontaneous mutations, and these mutations would occur at different times. Therefore, we set out to investigate the effect of deletions of MMR genes on antibiotic resistance in independently constructed strains through efficiency of plating assays (EOPs). We transduced Δ*mutL* and Δ*mutS* alleles into a Δ*yhdP* strain and assayed vancomycin resistance in five separate transductants. We observed the same level of resistance in all transductants (**Figure 1A**). The equal resistance phenotypes observed in all independent transductants demonstrate that loss of MMR is directly responsible for the resistance observed and secondary mutations are not necessary for the resistance.

**Figure 1.**
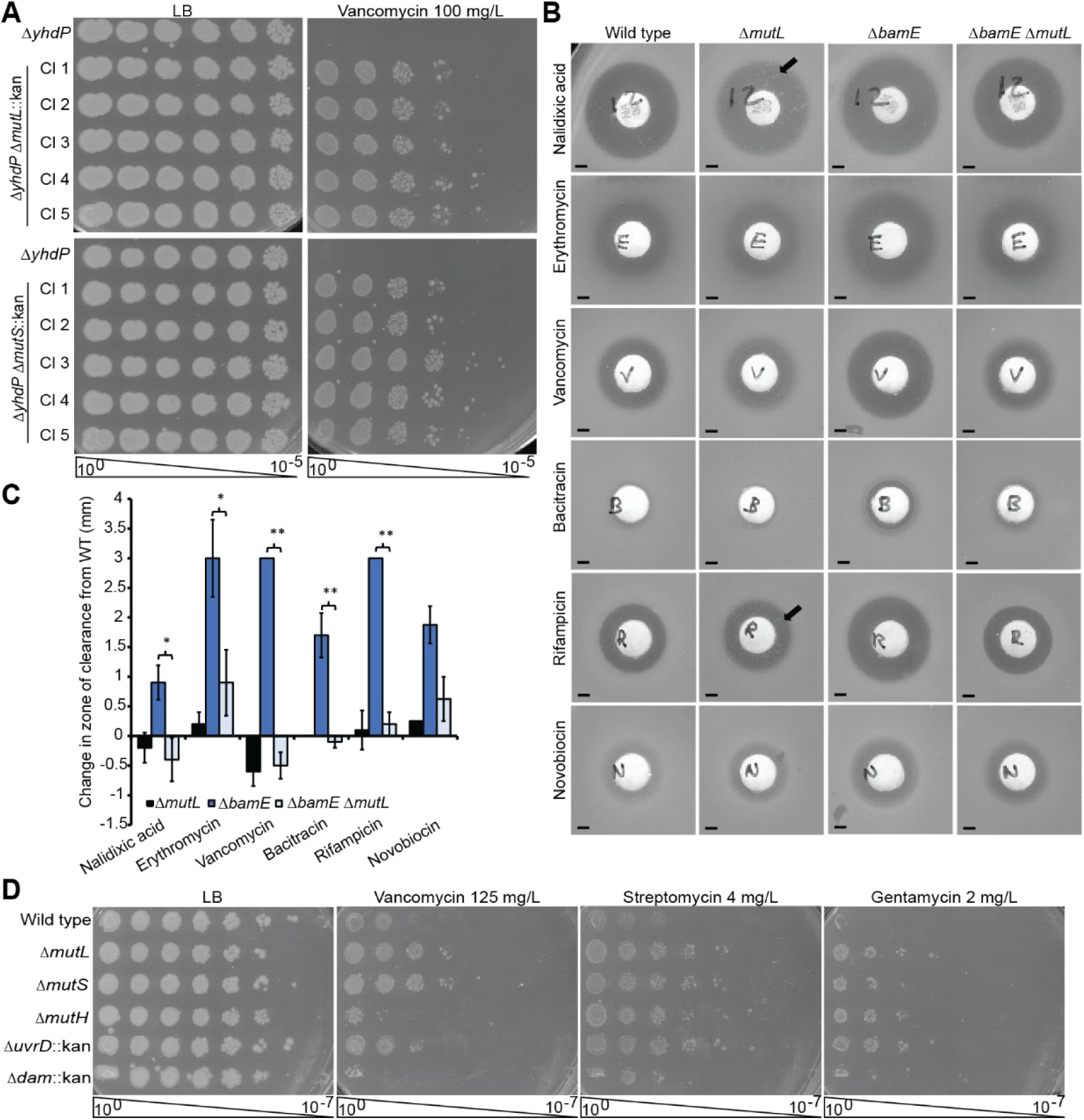
Increased antibiotic resistance occurs due to loss of MMR genes and not increased mutation rates. **(A)** Five transductants of *ΔmutL*::kan and *ΔmutS*::kan into a *ΔyhdP* strain were assayed and had comparable levels of vancomycin resistance as demonstrated by efficiency of plating assays (EOPs). **(B)** Kirby-Bauer tests show increased levels of antibiotic resistance in Δ*bamE ΔmutL* strains as compared to the Δ*bamE* strain as demonstrated by smaller zones of clearance. In contrast, black arrows indicate colonies that have acquired spontaneous resistance within the zone of clearance. Scale bars: 2 mm. **(C)** Quantification of Kirby-Bauer tests demonstrates increased resistance in the *ΔmutL ΔbamE* strain compared to the Δ*bamE* strain. Mean of five biological replicates ± SEM. * p<0.05 and ** p<0.01 by Student’s T test. **(D)** Loss of MMR genes at different steps in the MMR pathway confers varying degrees of antibiotic resistance. Inhibition of Δ*mutL* and Δ*mutS* causes higher levels of antibiotic resistance than wild type. Δ*mutH* is consistently less resistant than Δ*mutL* and Δ*mutS*, while the degree of resistance with Δ*uvrD* differs between antibiotics. However, Δ*dam*, which causes the same increase in mutations as loss of MMR, does not confer resistance. EOP images are representative of at least three independent experiments.

Next, we further investigated the resistance phenotype by performing Kirby-Bauer tests (i) to confirm the phenotype we observe is, in fact, resistance (i.e., an increase in MIC), (ii) to determine whether the resistance was vancomycin-specific or broader, and (iii) to investigate whether MMR mutants would confer resistance in OM-deficient mutants beyond Δ*yhdP*. We measured resistance based on the zone of clearance around the antibiotic disk: the smaller the zone of clearance, the more resistant the strain. Furthermore, this assay allows us to differentiate between changes in resistance of the strain from spontaneous resistance mutants occurring within the population (i.e., colonies within the zone of clearance) (**Figure 1B**, black arrows). Specifically, we assayed resistance in a strain with a deletion of *bamE*, a non-essential part of the β-barrel assembly machine (BAM) complex that inserts outer membrane proteins (OMPs) into the OM (31). This strain shows decreased resistance to several antibiotics that are generally excluded by the OM (**Figure 1BC**) (32). In the Δ*bamE* background, we observed significantly smaller zones of clearance with Δ*mutL* for several antibiotics to which *bamE* mutants have decreased resistance (**Figure 1BC**), demonstrating population-level increased resistance to the antibiotics. In addition, these data show that the increased resistance is not restricted to the Δ*yhdP* background or to vancomycin resistance.

We then asked if resistance occurs in a wild-type background and whether mutants in different steps of the MMR pathway cause equal levels of resistance. Our original screen detected insertions in *mutL*, *mutS* and *uvrD* as vancomycin resistant, but the screen was not saturating (26). We observed no bulk growth defects in strains with deletions of MMR genes (**Figure S2A**). We first performed Kirby-Bauer tests with a wide range of antibiotics to determine the antibiotics for which loss of MMR genes might cause a difference in resistance. Then, we performed EOPs and observed resistance to several antibiotics in a wild-type background with Δ*mutL* and Δ*mutS* that was very similar for each deletion (**Figure 1D**). We also confirmed that plasmid-based expression of *mutL* or *mutS* could restore sensitivity to their respective deletion strain (**Figure S2B**). When we assayed resistance to gentamicin and streptomycin by MIC assay in a wild-type background, we also observed a similar increase in MIC for the Δ*mutL* and Δ*mutS* strains (**Table S1**). These data confirm that loss of MMR genes can cause increased resistance in an otherwise wild-type strain background. In comparison to Δ*mutL* and Δ*mutS*, Δ*mutH* had an intermediate phenotype: *mutH* was equally or more resistant to antibiotics than the wild type, but less resistant than Δ*mutL* or Δ*mutS* strains (**Figure 1D**). Deletion of the helicase *uvrD* caused the strain to be more resistant than the Δ*mutH* strain, at times similar to the Δ*mutL* or Δ*mutS* strains. However, deletion of *dam*, encoding the Dam methylase responsible for the methylation that directs the strand specificity of MMR, had no effect on antibiotic resistance confirming that increased mutation rates are not sufficient for the resistance. Thus, although MMR mutations cause a direct increase in antibiotic resistance in a wild-type background, the level of resistance is not equal between the MMR mutants, suggesting that the method of MMR inhibition is important to the resistance. Altogether, we observed resistance in a wild-type background and in two different OM permeability mutants. We observed resistance to eight different antibiotics—vancomycin, streptomycin, gentamicin, rifampicin, nalidixic acid, bacitracin, novobiocin and erythromycin. These antibiotics do not share a class, molecular properties (e.g., size, charge), or mechanism of action, demonstrating the breath of this resistance.

### Loss of MMR increases antibiotic resistance in OM-deficient strains

As loss of each MMR pathway member leads to increased antibiotic resistance in a wild-type strain and resistance can be observed in a Δ*yhdP* strain, we next set out to determine whether loss of MMR could increase antibiotic resistance in other OM permeability mutants or whether the resistance was restricted to deficiencies in specific OM pathways. We identified strains with OM defects causing decreased resistance to antibiotics based on Kirby-Bauer tests. Then, we coupled deletion of the implicated genes with Δ*mutL* or Δ*mutS* and performed EOPs for antibiotics to which the OM mutants were less resistant. We began with BAM complex members (31), since the BAM complex has been implicated in OM permeability and serves as an antimicrobial target (33–36). BamE is a lipoprotein that serves a bridging function to coordinate activation of BamA and BamD, the essential members of the BAM complex (37). We observed that the D*bamE* strain showed decreased resistance to vancomycin, rifampicin, and bacitracin (**Figure 2A**). When D*bamE* was coupled with either Δ*mutL* or Δ*mutS,* we observed increased resistance in the double mutants. This level of resistance was much greater than the Δ*bamE* parent strain but less than that of wild type. To confirm that the resistance was also independent from secondary mutations in the Δ*bamE* background, we made three individual double mutants and observed similar resistance in each of them (**Figure 2A**). We observed similar results in Δ*bamB* (**Figure 2B**), which codes for another non-essential BAM lipoprotein (38, 39), and *bamA101* (**Figure 2C**), a promoter-down mutant of the gene coding for the essential BamA OMP (40), backgrounds.

**Fig 2.**
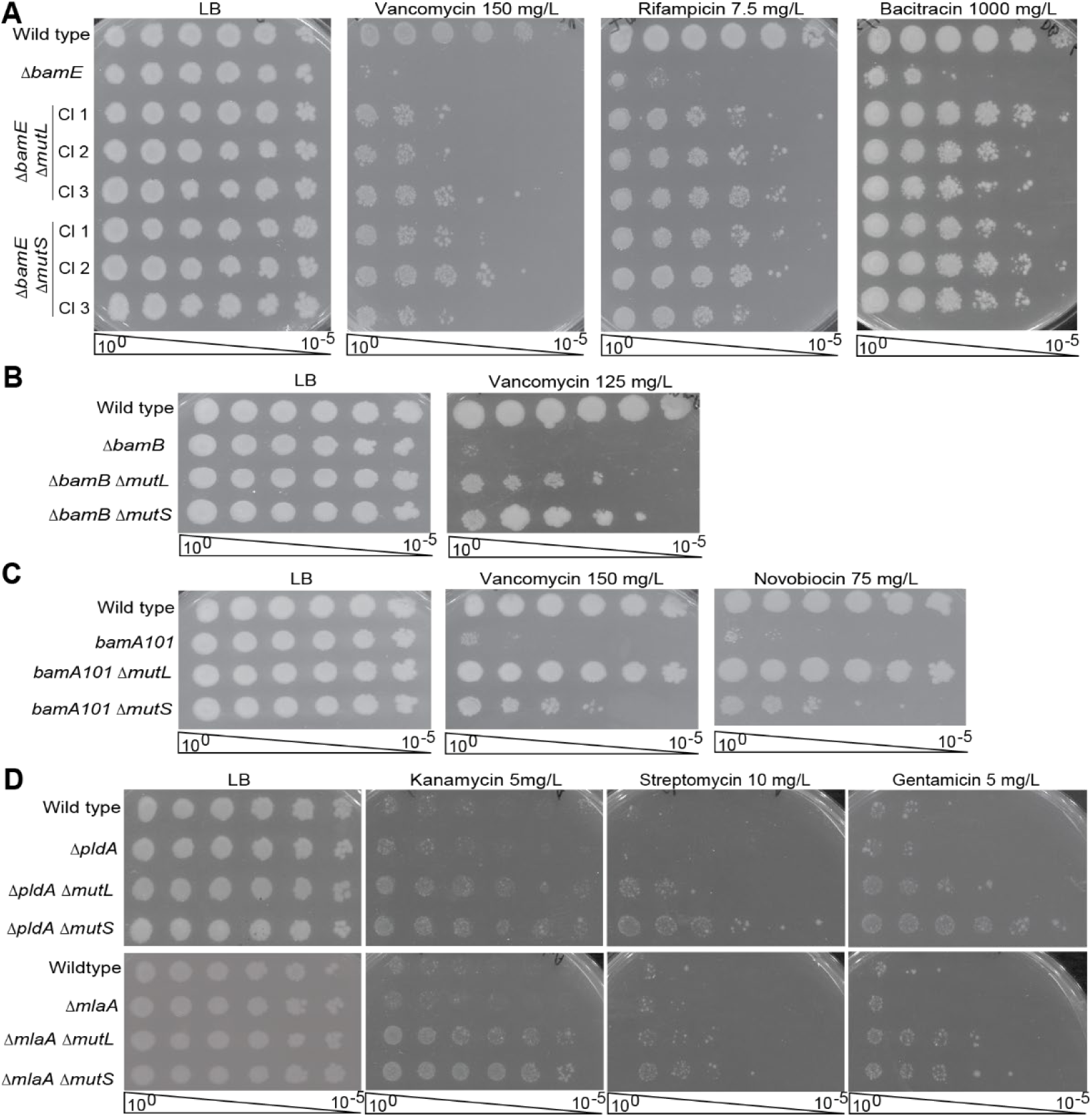
Loss of MMR genes can overcome antibiotic sensitivity in strains with a compromised outer membrane. Loss of MMR genes can increase antibiotic resistance, overcoming OM deficiencies in multiple mutants in BAM (β barrel assembly machinery). BamB and BamE are non-essential BAM lipoproteins, while BamA is the essential OMP that facilitates OM insertion of newly synthesized OMPs. **(A)** Δ*bamE* causes decreased resistance to the indicated antibiotics. Combining Δ*mutL* or Δ*mutS* with *ΔbamE* restores partial resistance to these antibiotics. Three separate transductants are shown. Strains with Δ*bamB* **(B)** and lowered expression of *bamA* (*bamA101*) **(C)** have less resistance to vancomycin, and *bamA101* has less resistance to novobiocin. Δ*mutL* or Δ*mutS* partially or fully restores resistance to wild type levels. **(D)** PldA is a phospholipase that cleaves phospholipids in the outer leaflet of the OM while, MlaA is the OM component of a systems that returns mislocalized phospholipids from the outer leaflet of the OM to the IM. Loss of MMR genes in Δ*pldA* and Δ*mlaA* increases antibiotic resistance above that of wild type. Images are representative of three independent experiments.

To investigate other OM functions, we constructed Δ*mlaA* and Δ*pldA* mutants coupled with deletion of MMR genes. PldA is a phospholipase, which contributes to the asymmetry of the OM by cleaving phospholipids that have become mislocalized to the outer leaflet of the outer membrane and initiating a pathway that leads to upregulation of LPS production (41–45). MlaA is an outer membrane lipoprotein involved in a retrograde phospholipid trafficking pathway that maintains outer membrane lipid asymmetry by removing mislocalized outer leaflet phospholipids and transporting them back to the IM (46–48). Once again, we observed resistance with either Δ*mutL* or Δ*mutS* when compared to the parent OM mutants, although these OM mutants were equally resistant to the wild-type strain (**Figure 2D**). Our data demonstrate that the phenotype of resistance seen with loss of MMR genes can overcome multiple kinds of increased OM permeability. As resistance can be increased in OM mutants in differing pathways (i.e., phospholipid transport, OMP biogenesis, OM asymmetry), these data demonstrate that the mechanism of increased resistance is not tied to a specific OM biosynthesis pathway, suggesting the resistance may not relate to strengthening of the OM.

### Increased antibiotic resistance via loss of MMR extends beyond *E. coli*

MMR is an evolutionarily conserved process across many organisms (14). As we characterized the scope of the resistance caused by loss of MMR genes in *E. coli* K-12, we turned to whether the resistance extended beyond *E. coli* or whether it was *E. coli* specific. Therefore, we investigated whether loss of MMR genes caused increased antibiotic resistance in *Salmonella enterica* serovar Typhimurium LT2, the type strain for the serovar. Using λ Red recombineering, we recombineered a kanamycin resistance cassette into the chromosome in place of the *mutL* and *mutS* genes (49). We then identified possible changes in antibiotic resistance by Kirby-Bauer tests and tested the effect of the Δ*mutL* and Δ*mutS* mutations in these strains by EOP. The newly constructed Δ*mutL* and Δ*mutS* strains showed increased antibiotic resistance when compared to wild-type *S.* Typhimurium (**Figure 3**). The antibiotics for which we observed increased resistance, and the level of resistance observed, were similar to that which we observed with *E. coli* K-12. These data demonstrate loss of MMR genes leads to antibiotic resistance beyond *E. coli* K-12 and suggests that the resistance is due to the same mechanism, since the pattern of resistance is similar between the species.

**Figure 3.**
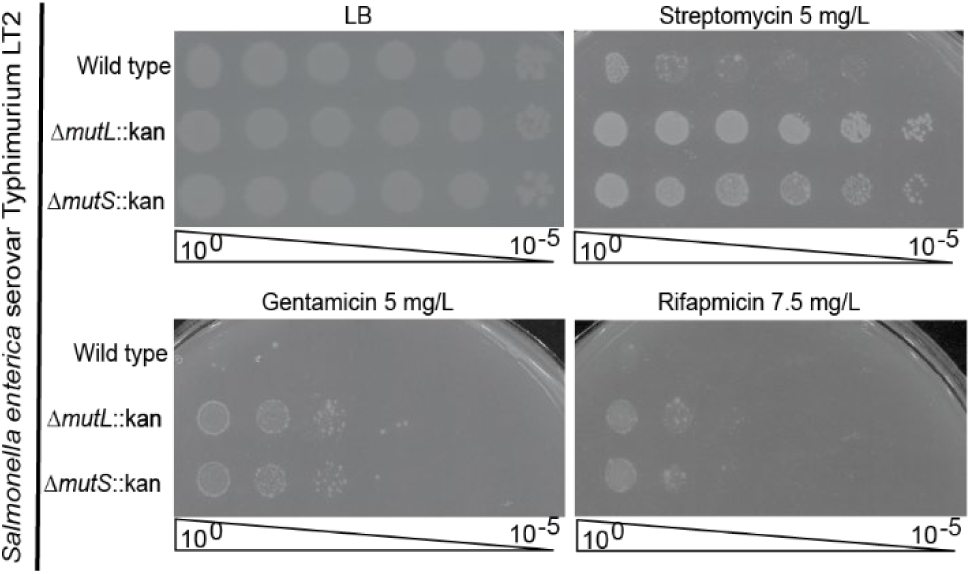
Increased antibiotic resistance from loss of MMR genes extends beyond *E. coli*. Deletion of *mutL* or *mutS* in *Salmonella enterica* Serovar Typhimurium LT2 results in increased antibiotic resistance compared to wild type. Thus, loss of MMR genes as a mechanism for antibiotic resistance is not specific to *E. coli*. EOP images are representative of three independent experiments.

### Reported rates of homoeologous recombination correlate with increased antibiotic resistance

Our work thus far has established that loss of MMR genes leads to increased antibiotic resistance which can overcome increased OM permeability. We then turned our attention from characterizing the scope of resistance to investigating possible mechanisms of the resistance in *E. coli*. Since MMR can repair DNA damage beyond mismatches, the resistance we observe could occur through activation of the SOS response, *E. coli’s* cellular stress response to severe DNA damage. The SOS response is activated when the cell senses an accumulation of single-stranded DNA (50). The presence of RecA bound to single-stranded DNA causes the self-cleavage of LexA (the SOS transcriptional repressor) which removes LexA from the SOS boxes in the promoters of regulated genes, derepressing transcription (51–53). We thus investigated the SOS response’s role in the increased antibiotic resistance caused by loss of MMR genes. First, we determined whether increased resistance from loss of MMR genes could occur in the absence of *recA*, which binds to the single-stranded DNA. Δ*recA* caused decreased resistance to vancomycin when compared to wild type and we observed a similar level of resistance between the Δ*recA* and Δ*mutL* Δ*recA* strains (**Figure 4A**). However, Δ*recA* in a Δ*yhdP* background caused resistance to increase to wild-type levels, higher than even Δ*yhdP* Δ*mutL.* Interestingly, when Δ*recA* was coupled with Δ*yhdP* Δ*mutL*, the strain showed a similar loss of resistance to a Δ*yhdP* strain, much less than either of the double mutants. These complex genetic interactions, especially the effects of RecA on antibiotic resistance, made it difficult to assess the role of SOS in the MMR mediated resistance pathway. This is especially true as RecA functions outside of the SOS response and is necessary for homologous recombination, so loss of RecA could result in phenotypes unrelated to SOS inactivation.

**Figure 4.**
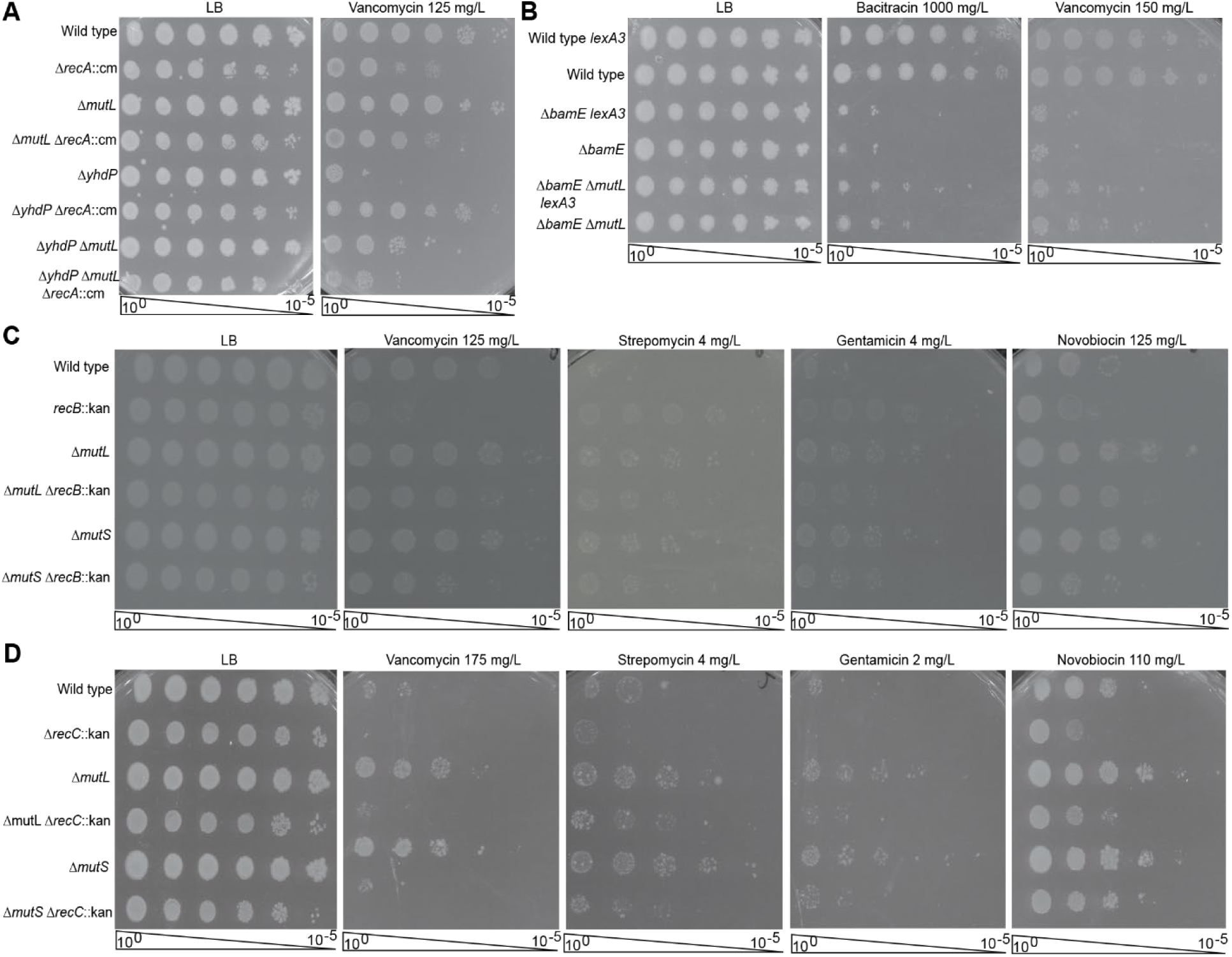
Increased antibiotic resistance with loss of MMR genes is independent SOS and is linked to homologous recombination. The SOS response is *E. coli*’s response to DNA damage that leads to exposure of single-stranded DNA. **(A)** RecA is required for both homologous recombination and the SOS response. RecA has complex genetic interactions with antibiotic resistance mediated by loss of MMR genes. Δ*recA* causes increased resistance to vancomycin in wild-type and Δ*mutL* backgrounds. In a Δ*yhdP* background, Δ*recA* causes increased vancomycin resistance; however, resistance in Δ*mutL* Δ*yhdP* Δ*recA* is similar to that of Δ*yhdP* alone. **(B)** Inactivation of the SOS response via *lexA3* (a non-cleavable mutant of the SOS transcriptional regulator) does not change antibiotic resistance mediated by loss of MMR genes. **(C-D)** Inhibition of MMR has previously been linked to increased homologous recombination rates. Δ*recB* (C) and Δ*recC* (D), which reduce homologous recombination rates, result in decreased antibiotic resistance in both the wild type and MMR gene deletion backgrounds for most of the assayed antibiotics. EOP images are representative of three independent experiments.

To inactivate the SOS response by preventing the release of the SOS repressor LexA from SOS boxes, we transduced a non-cleavable *lexA* allele (54) into our Δ*bamE*, Δ*bamE* Δ*mutL*, and Δ*bamE* Δ*mutS* strains. The increased resistance from Δ*mutL* and Δ*mutS* was not changed when the non-inducible *lexA* allele was introduced (**Figure 4B**), demonstrating that the SOS response does not play a role in the pathway linking loss of MMR genes to increased levels of antibiotic resistance. This is consistent with the lack of similar phenotypes caused by deletion of genes in other DNA repair pathways (**Figure S3**). We also tested whether several other stress responses (see **Figure S4** for examples) were necessary for the increased resistance induced by loss of MMR genes and did not find any links to these pathways.

The phenotypes of the Δ*recA* mutants with loss of MMR genes suggested RecA could be playing a role in the resistance phenotypes we observe with loss of MMR genes. The other major role of RecA is in homologous recombination (55). The RecA effect on increased antibiotic resistance with loss of MMR genes brought to mind the effect of MMR on homoeologous recombination, homologous recombination between divergent (non-identical) DNA strands (56). It has been established that inhibition of MMR leads to increased rates of homoeologous recombination as MutS and MutL can block RecA-mediated strand exchange when nucleotide mismatches occur (56–58). However, the change in rates of recombination is not equal for all MMR genes. Inhibition of MMR via Δ*mutS* or Δ*mutL* increases recombination rates by 735-fold, while Δ*mutH* and Δ*uvrD* increase rates 22-fold and 5-fold, respectively, with simultaneous inhibition of MutH and UvrD producing similar phenotypes to loss of MutL or MutS (56–58). Interestingly, aside from UvrD, which also functions in nucleotide excision repair and so may have secondary effects (59), the pattern of increased rates of homoeologous recombination is similar to the pattern of increased antibiotic we observed with the deletions of the various MMR genes (**Figure 1D**, **S5**). This, combined with the effect of RecA on antibiotic resistance in the Δ*mutL* and Δ*yhdP* Δ*mutL* backgrounds, caused us to ask whether homologous recombination was involved in increased antibiotic resistance with loss of MMR genes (60).

Thus, we combined deletions of MMR genes with deletions of genes in the main homologous recombination pathway in *E. coli*, the RecBCD pathway (61). If the increase in recombination rates in strains with MMR gene deletions is important for antibiotic resistance, then strains with reduced recombination rates should have lower resistance to antibiotics. Therefore, we combined deletions of *recB* and *recC*, which code for proteins forming a complex with endonuclease and helicase activity that initiates recombination in the RecBCD pathway (61), with Δ*mutL* or Δ*mutS* and assayed antibiotic resistance. Δ*recB* or Δ*recC* strains have an approximately 100-fold lower rate of chromosomal recombination compared to wild type (62–67). We observed that while the Δ*mutL* or Δ*mutS* single mutants displayed increased resistance compared to the wild type, the Δ*recB* and Δ*recC* single mutants were generally less resistant than wild type to the same antibiotics (**Figure 4CD**). When we coupled Δ*mutL* or Δ*mutS* with Δ*recB* or Δ*recC*, we observed a similar lack of resistance as the recombination mutants alone, demonstrating that loss of MMR genes does not confer increased resistance in a recombination deficient background. These results are similar to those we observed with loss of RecA (**Figure 4A**). However, the effects of Δ*recB* and Δ*recC* are not universal—loss of these genes does not appear to cause a large effect on streptomycin resistance (**Figure CD**). Together, these data demonstrate that homologous and/or homoeologous recombination is important for antibiotic resistance and suggest that the increase in resistance to several antibiotics caused by loss of MMR genes is due to increased recombination rates.

### Loss of MMR genes leads to cell lysis

Changes in antibiotic resistance similar to what we have observed with MMR mutants can be caused by physiological changes to the OM (7, 9–12), which serves as an intrinsic barrier against antibiotic entry (12, 68, 69). Although our data did not tie the increased resistance observed with loss of MMR genes to a specific OM biogenesis pathway, we used chlorophenyl red-β-D-galactopyranoside (CPRG) to test OM permeability and cell integrity in *E. coli*, as has been previously described (70). When CPRG contacts β-galactosidase either in the cytoplasm or the extracellular environment, the bond between chlorophenyl red and galactose is cleaved, causing a color change from yellow to red. We grew our MMR gene deletion strains in the presence of CPRG and IPTG, to induce *lac* operon expression, and assayed the ability of CPRG and β-galactosidase to come into contact. The Δe*lyC* strain served as a positive control as this strain has been shown to lyse at low temperatures (70). The MMR gene deletion strains all showed increased CPRG activity when compared to the wild-type strain (**Figure 5A**, **S6A**). Moreover, OM deficient strains, such as Δ*bamE*, showed increased CPRG activity as expected. However, when Δ*bamE* was coupled with the Δ*mutL* or Δ*mutS*, the strains demonstrated even more CPRG activity. These data demonstrate either the MMR gene deletion strains are more permeable to CPRG than wild type or CPRG is more able to contact β-galactosidase because β-galactosidase is escaping the cell (i.e., through lysis). We decided to investigate whether the strains with deletions of MMR genes were exhibiting some level of lysis. We performed western blots on the precipitated supernatant from overnight cultures to detect the presence of the cytoplasmic chaperonin protein GroEL in the culture media. We observed higher GroEL levels in the culture supernatant the MMR gene deletions when compared to the wild type (**Figure 5B**). When we quantitated GroEL levels, we found they were 6- to 14-fold higher in the MMR mutants than in wild type (**Figure S6B**).

**Figure 5.**
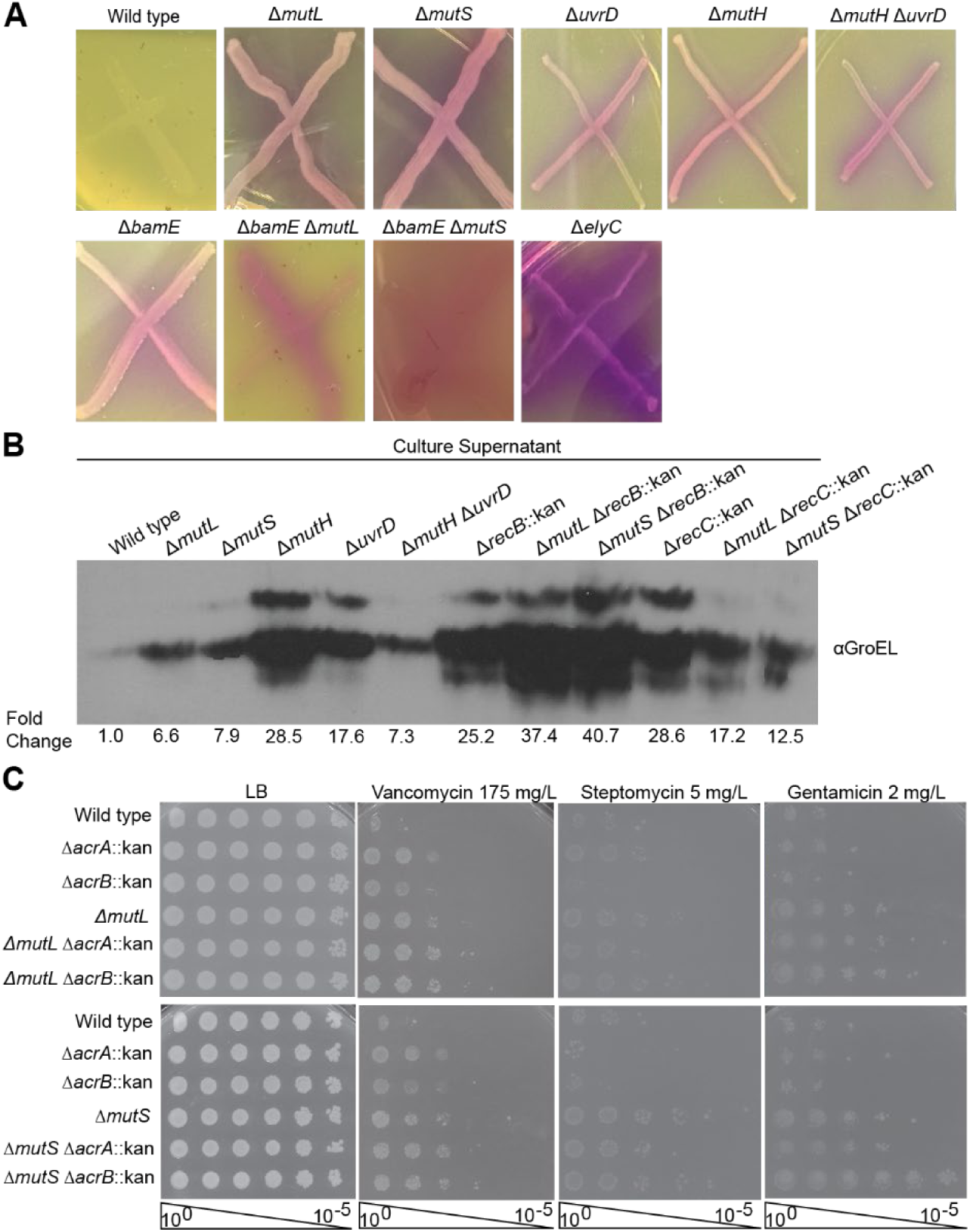
Loss of MMR genes causes increased cell lysis. **(A)** CPRG is a poorly cell-permeable, colorimetric β-galactosidase substrate that must encounter β-galactosidase, either through increased OM permeability or cell lysis, to be cleaved releasing chlorophenol red. Deletion of MMR genes causes increased CPRG cleavage, suggesting either increased OM permeability or cell lysis. **(B)** To assay lysis, immunoblots detecting the cytoplasmic protein GroEL were performed on the TCA precipitated supernatant of overnight cultures. Strains with deletions of MMR genes show higher levels of GroEL in the supernatant than wild-type cells do, demonstrating increased lysis. However, this lysis was not decreased by loss of RecB or RecC. **(C)** Necrosignaling may play a role in resistance to some assayed antibiotics. AcrA released from lysing cells is sensed by neighboring cells to induce a decrease in antibiotic resistance (71, 72). The change in resistance from loss of necrosignaling (Δ*acrA*) was compared to that of loss of the AcrAB-TolC efflux pump (Δ*acrB*). While Δ*acrA* did not change levels of vancomycin or gentamicin resistance in the Δ*mutL* and Δ*mutS* backgrounds, Δ*acrA* resulted in some loss of resistance to streptomycin. All images are representative of at least three independent experiments.

We wondered whether the increased lysis rates were due to illegitimate homologous recombination and so tested lysis in strains with Δ*mutL* or Δ*mutS* combined with Δ*recB* and Δ*recC*. However, the Δ*recB* and Δ*recC* strains showed higher levels of lysis than Δ*mutL* or Δ*mutS* caused, complicating analysis (**Figure 5B**, **S6B**). Nevertheless, Δ*mutL* or Δ*mutS* did not cause any additional increase in lysis in the Δ*recB* and Δ*recC* strains, suggesting lysis in the MMR gene deletion strains strains may be linked to increased recombination. As the double mutant strains are more sensitive to antibiotics than strains lacking MMR genes alone, these data also suggest that, if lysis is involved in antibiotic resistance, lysis is not sufficient, or a specific level of lysis is necessary to promote resistance.

We further wondered whether the antibiotic resistance phenotypes we observed might be due to the phenomenon of “necrosignaling” recently described in swarming *E.* coli (71, 72). This pathway relies on AcrA, the lipoprotein component of the AcrAB-TolC RND family efflux pump (73, 74). Dead cells release a necrosignal in the form of AcrA which interacts with the OM of the population of the swarm that has yet to encounter the antibiotic. The resistance of the signaled cells to the antibiotic is increased due to the upregulation of efflux pumps and metabolic changes to the cells. We constructed Δ*mutL* or Δ*mutS* coupled with Δ*acrA* and Δ*acrB* and investigated whether these mutations would affect antibiotic resistance. Changes in resistance based on efflux should be equal between the Δ*acrA* and Δ*acrB* mutants, while necrosignaling would be affected more strongly by Δ*acrA*. Resistance to vancomycin, streptomycin, and gentamicin were not strongly affected by the loss of AcrB (**Figure 5C**). Δ*acrA* did not change vancomycin or gentamicin resistance in the MMR deficient backgrounds; however, streptomycin resistance was reduced to near wild-type levels in the MMR Δ*acrA* double mutants (**Figure 5C**). Interestingly, this is the same antibiotic for which we did not observe a change in the resistance conferred by MMR gene deletions in backgrounds with decrease in recombination. Therefore, necrosignaling may be playing a role in increasing resistance to some antibiotics in MMR mutants, while rates of recombination are important for others. These data demonstrate that loss of MMR causes some level of cell lysis within the cell population. Further, this lysis may act to affect resistance to some antibiotics through a necrosignaling pathway.

## Discussion

Here, we present evidence that a novel pathway acts to increase population-wide antibiotic resistance in *E. coli* and *S.* Typhimurium when MMR is inhibited through deletion of MMR genes. This pathway acts on antibiotics of many classes, sizes, and mechanisms of action; moreover, it is independent of increased mutation rates. This increase in resistance can be observed in wild-type strains as well as in various mutants with increased OM permeability. Increased mutations rates in hypermutator strains such as MMR mutants have been shown to be associated with antibiotic resistance (30). The pathway we have described provides a mechanism by which MMR-inhibited hypermutator strains could survive in the presence of antibiotics, giving them time to develop mutations causing true resistance. This also suggests that the proportion of MMR mutants in a population may start to expand under antibiotic treatment due to their intrinsic resistance, providing increased opportunity for antibiotic specific mutations conferring high-level resistance to occur. Previous work has shown mutations causing low-level resistance can lead to a greater chance of development of further resistance mutations, allowing strains to cross the clinical breakpoint into being resistant (7, 9–12). Therefore, this pathway mediated by loss of MMR genes, and the low-level resistance it causes, could be an important contributing factor to the development and spread of resistance alleles.

The pathway we have described causes an increase in resistance to a broad range of antibiotics via loss of MMR genes, independent of the strains’ increased mutation rates. We can confidently say the resistance is independent of mutation rates because we were able to transduce MMR gene deletions into multiple strains and backgrounds and observe a similar increase in resistance. Furthermore, we can differentiate between resistance mutations occurring within a population and a strain with increased resistance through Kirby-Bauer assays where population-level resistance determines the zone of clearance around the disc, while spontaneous mutants within the population appear as colonies within the zone of clearance. Finally, loss of the Dam methylase has the same effect on mutation rates as loss of MMR genes but does not confer increased antibiotic resistance, demonstrating that increased mutation rates are not sufficient for increased resistance. In addition to this robust evidence, the increased mutation rates should not be sufficient to cause a consistent increase in resistance between many independent strains. For example, the rate of spontaneous base-pair substitutions of *E. coli* K-12 is 2 × 10^-10^ mutations per nucleotide per generation (75, 76). When MMR is inactivated, there will be a 15-20-fold increase in mutations due to the activity of repair pathways mechanisms of repair that cannot differentiate between daughter and parental strands of DNA (77), corresponding to approximately 1 mutation per 75 generations of growth (76, 77). Given our attempts to avoid extended culture time (i.e., minimizing time in stationary phase) and passaging of these strains (i.e., by streaking fresh plates from glycerol stocks immediately before each experiment), this mutation rate would be insufficient for resistance mutations to consistently occur and spread throughout the cell population before resistance is assayed.

That the pathway is independent of increased mutation rates opens new questions about the relationship between DNA repair, mutation rates, and antibiotic resistance, specifically the mechanism linking loss of MMR genes to increased antibiotic resistance. Our data provide some insights into this mechanism, although elucidating the full mechanism is likely to be many years of work. Our data links the increased resistance to several antibiotics to increased recombination rates and suggest that some level of cell lysis within the population may be involved in the resistance to others. Without MMR genes, cells will have higher rates of homoeologous recombination, where sequences between recombining strands differ. Thus, there will be a higher probability of illegitimate or unsuccessful recombination. We believe that this may lead to low levels of lysis within the population that, in turn, increase antibiotic resistance. This increased resistance could be through necrosignaling causing physiological changes in surviving cells (71, 72), as we have demonstrated contributes to changes in streptomycin resistance. For other antibiotics where necrosignaling does not appear to play a role, antibiotics could be exposed to and bind their targets outside of cells lowering their effective concentration. In this way, interactions between cell debris and antibiotics could lower the amount of the antibiotic that can enter living cells. One example of this type of extracellular interaction of an antibiotic with its target leading to resistance comes from a spontaneous vancomycin resistance mutation in *E. coli* that causes it to attach peptidoglycan subunits to its LPS (78). Vancomycin interacts with the cell surface instead of penetrating the OM, allowing the mutant to grow on much higher concentrations of vancomycin than the wild type.

It is also possible that illegitimate or unsuccessful recombination increases antibiotic resistance independently from the increased lysis we observe in strains with deletions of MMR genes. For instance, these events may initiate a signaling cascade that causes physiological changes increasing the cells’ antibiotic resistance. Transcriptional changes leading to multidrug resistance have been associated with bacterial stress responses including those mediated by MarA, SoxS, and SdiA (79, 80). We previously conducted RNA sequencing of untreated MMR mutants, but did not resolve any transcriptional changes that were required for changes to antibiotic resistance. However, it may be that, given the sporadic nature of recombination events, single cell RNA-Seq will be required for investigating this possibility as this technology improves for bacterial applications. In addition, it may be that the illegitimate recombination events inducing transcriptional changes in MMR mutants occur specifically after antibiotic treatment. We are currently investigating this possibility. Any mediating steps we uncover in this pathway would be potential targets for decreasing the development of resistance mutations. Finally, it is possible that antibiotic treatment induces recombination events that directly increase resistance (e.g., through gene duplication events); however, this mechanism would not repeatedly produce equal levels of antibiotic resistance and so we consider this a less likely possibility.

Much like MMR, the links proposed between recombination and antibiotic resistance have mainly been related to the spread of resistance alleles through pathways that require recombination (e.g., conjugation or transduction) (55). More recently, DNA repair via homologous recombination has been linked to persistence after fluoroquinolone treatment (81). Our data show that the effects of recombination rates on antibiotic resistance can extend beyond antibiotics that target DNA replication or directly cause double-strand DNA breaks. This suggests that recombination may be playing a larger role in antibiotic responses—perhaps through repair of DNA damage that occurs due to metabolic changes during antibiotic treatment (82, 83). It has been well studied that accumulation of DNA damage can lead to stress response activity (51, 84–86). It may be that DNA damage occurring during antibiotic treatment works in conjunction with increased homoeologous recombination to activate further resistance. These are intriguing areas for further investigation.

We have shown a promising link between decreased rates of recombination and decreased resistance to antibiotics, and a similar mechanism may extend beyond *E. coli*. Thus far, our mechanistic studies have been performed in *E. coli.* However, there is some evidence that loss of MMR genes leads to increased rates of recombination in *S.* Typhimurium (56, 58). MMR is conserved within eukaryotic and prokaryotic systems (87, 88). More importantly, the pattern of suppression of homoeologous recombination via MMR is also conserved through prokaryotic and eukaryotic systems, meaning loss of MMR genes, regardless of system, could result in stress responses and adaptation to stress. For instance, studies in yeast have revealed that suppression of homoeologous recombination occurs via a mechanism that involves RecQ family helicase SGS1 and MutSα (89, 90). Elucidating this pathway further will allow for a more detailed perspective on how increased antibiotic resistance and possibly resistance to other stresses occurs with MMR mutants in different organisms.

## Materials and Methods

### Bacterial strains and growth conditions

All strains used in this study are listed in **Table S2**. Cultures were grown in LB Lennox media at 37°C. Knockout strains in *E. coli* K-12 MG1655 were constructed with Keio collection alleles (91) using P1*vir* transduction (92). The pCP20 plasmid, which encodes a FLP recombinase-FRT system to remove antibiotic resistance cassettes, was used to remove resistance cassettes after transduction (49). New alleles in *S.* Typhimurium and *E. coli* were constructed with λ-Red recombineering using the pKD46 plasmid using the indicated primers (**Table S3**) and the kanR cassette from pKD13 or the cmR cassette from pKD3 as a template, as has been described (49). In *S.* Typhimurium, the recombineering plasmid was cleared by growth at 37 °C. For *E. coli* strains, newly constructed alleles were moved to clean backgrounds using P1*vir* transduction.

Complementation plasmids for *mutL* and *mutS* were constructed by using the mutL_fwd/rev and mutS_fwd/rev primers (**Table S3**) to amplify *mutL* or *mutS*, respectively, from genomic DNA. pBAD33 (93) was amplified using the pBAD-fwd and pBAD-rev primers and the fragments were combined with HiFi Assembly (New England Biolabs) as per the manufacturer’s instructions. Before experiments, strains were streak for single colonies from glycerol stocks and individual colonies cultured immediately for experiments. MMR mutant strains were not stored after growth or passaged. When necessary for plasmid maintenance, cultures were supplemented with 25 μg/mL kanamycin (Gold Biotechnology) or 20 μg/mL chloramphenicol (Gold Biotechnology). For assay of resistance phenotypes, LB was supplemented with the indicated concentrations of vancomycin (Gold Biotechnology), streptomycin (Gold Biotechnology), gentamicin (Gold Biotechnology), rifampicin (Gold Biotechnology), bacitracin (Gold Biotechnology), novobiocin (Gold Biotechnology), or kanamycin (Gold Biotechnology).

### Minimum inhibitory concentration assay

Strains were grown overnight in LB. Then, cultures were normalized to an OD of 0.1 and further diluted 1:1000 in LB. Next, cultures were pipetted into a 96-well plate at a volume of 98uL. Using a multichannel pipette, 2uL of 1.5-fold antibiotic dilutions were added to the cultures. Strains were incubated at 37°C overnight. The next day, the OD_600_ was read using a BioTek Synergy H1 plate reader. The lowest concentration where growth is inhibited was considered to be the MIC. The geometric mean of the MICs from three independent experiments was calculated.

### Efficiency of plating Assays (EOP)

Cultures were grown overnight in LB. 200 µL of overnight culture were pipetted into 96-well plates then serial diluted by a factor of 10. Dilutions were then plated onto the appropriate plates using a 48 well pin tool and the plates incubated overnight at 37°C.

### Kirby-Bauer Test

Strains were grown overnight. After overnight growth, 100 µL of culture was diluted into 3 mL of melted LB top agar and then the mixture was poured onto a LB agar plate until the top agar fully covered the plate. Once solidified, antibiotic discs (BD BBL™ Sensi-Disc™) were placed on the plate and the plate was incubated overnight at 37°C. For antibiotics that require higher amounts than provided by commercial discs, antibiotics in the following amounts were added to blank discs and dried before placing on the plate: vancomycin 300 µg, erythromycin 150 µg, novobiocin 200 µg, bacitracin 1 mg, rifampicin 20 µg. The following day, zones of inhibition were measured in 3 different directions and averaged.

### CPRG Assay

CPRG assays were performed as previously described with minor variations (27, 70). LB agar plates were supplemented with 20 µg/mL of CPRG and 50 µM IPTG. For plate assays, bacteria were streaked out onto a CPRG LB agar plate and let to grow overnight in the dark at room temperature. For quantification, liquid CPRG assays were performed by growing strains overnight at 37 °C with 50 µM IPTG. 200 µL of cells were spun down at 3700 rpm for 2 minutes in a 96-well plate. 100 µL of supernatant were transferred into a new 96-well plate, 100 µL of CPRG were then added and plate was incubated at 37 °C for 1 hour. Chlorophenol red absorption was read at a 575 nm wavelength.

### Immunoblot analysis for lysis via GroEL detection

The trichloroacetic acid (TCA) precipitation protocol was adapted from Ruiz, *et al.* (28). Cultures were grown overnight in LB at 37 ⁰C. After growth, 1.2 mL of culture was spun down at room temperature, 16,000 x g for 2 minutes. The supernatant was filtered through a 0.22 µm syringe filter, then 900 µL of filtered supernatant was transferred to a new microcentrifuge tube and proteins were precipitated with 100 µL of 100% TCA. TCA-precipitated proteins were then resuspended in 50 µl 1XSDS loading buffer. 20 µL of sample was loaded onto a 12% SDS-PAGE gel and the gel was run for 1 hour at 115 volts. Proteins from the gel were transferred to a nitrocellulose membrane. The GroEL chaperonin was detected with a GroEL primary antibody (Sigma) at a 1:30,000 dilution. The secondary antibody was goat anti-rabbit conjugated to horseradish peroxidase (Prometheus) at a 1:100,000 dilution. Detection of the signal was completed by using Prosignal Pico ECL (Prometheus) and Prosignal enhanced chemiluminescence (ECL) blotting film (Prometheus). Quantification of proteins levels was performed with ImageJ.

## Acknowledgments

We thank the members of the Mitchell lab, Jolene Ramsey and her Lab (Texas A&M University), Deborah Siegele (Texas A&M University), James Smith (Texas A&M University), Jonathan Sczepanski (Texas A&M University), Natividad Ruiz (Ohio State University), and Anna Konovalova (UT Health Houston) for feedback, constructive criticism, and productive discussions of this project. We also thank Deborah Siegele for the *lexA3* allele and James Slauch (University of Illinois Urbana-Champaign) for the wild-type *S.* Typhimurium strain. This work was supported by the National Institute of Allergy and Infectious Diseases under grants R01-AI155915 and R03-AI177677 (to A.M.M.) and by Texas A&M University startup funds.

