## Supplemental Figures and Tables for "Loss of DNA mismatch repair genes leads to acquisition of antibiotic resistance independent of secondary mutations"

**This file includes:**

Figures S1 to S6
Tables S1 to S3
Supplemental References

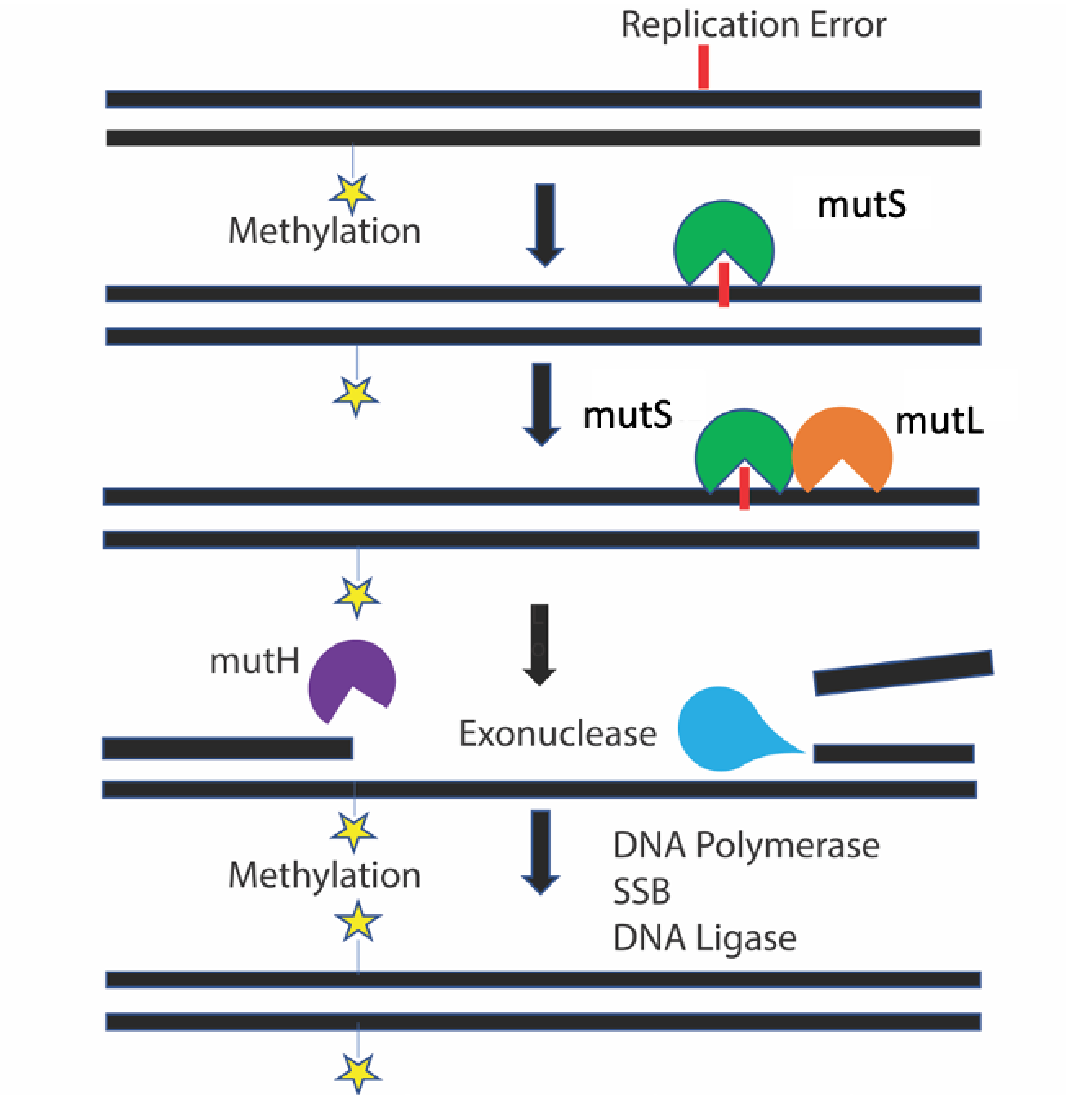


**Fig. S1. DNA Mismatch Repair Pathway.** A nucleotide mismatch formed during DNA replication is first recognized by MutS. MutS then recruits MutL and forms a heterodimer complex at the site of the mismatch. This complex recruits MutH which scans to the nearest GATC site and makes a nick on the unmethylated strand. UvrD unwinds the DNA allowing the exonuclease to enter and degrade the DNA strand past the point of the mismatch with single stranded binding protein (SSB) stabilizing the DNA. DNA polymerase resynthesizes the DNA strand and DNA ligase ligates the remaining nick. Finally, DAM methylates the nascent DNA strand.

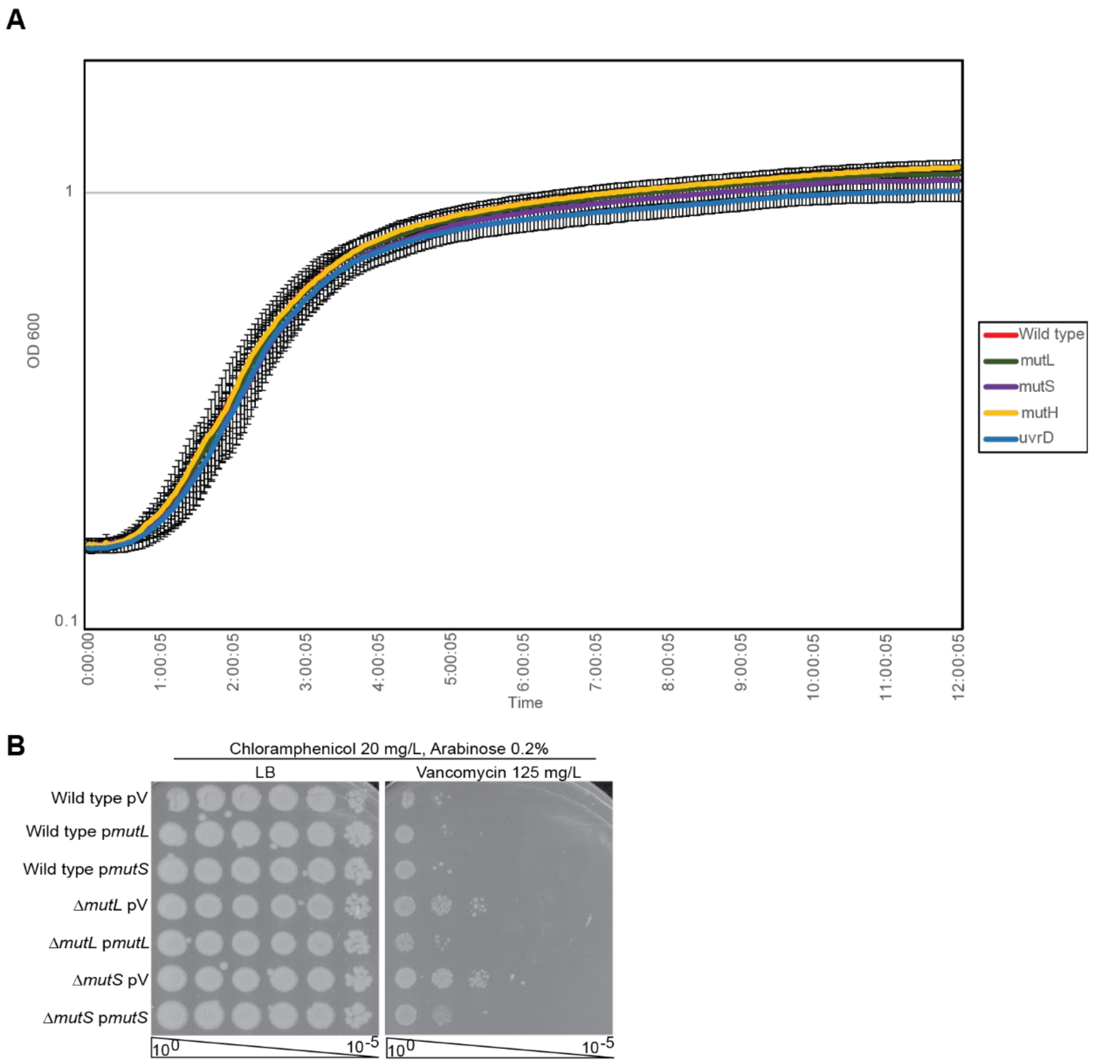


**Fig. S2. Loss of MMR does not cause a growth defect. (A)** Strains with deletions in MMR genes and wild-type cells were grown 37 °C for 12 hours. No growth differences between the strains were observed. Data are representative of 3 independent experiments. Mean ±Std Dev is shown. **(B)** Complementation of MMR gene deletions returns antibiotic resistance to wild-type levels. Plasmid-based copies of *mutL* or *mutS* were expressed from the arabinose inducible $P_{BAD}$ promoter in the indicated strain backgrounds.

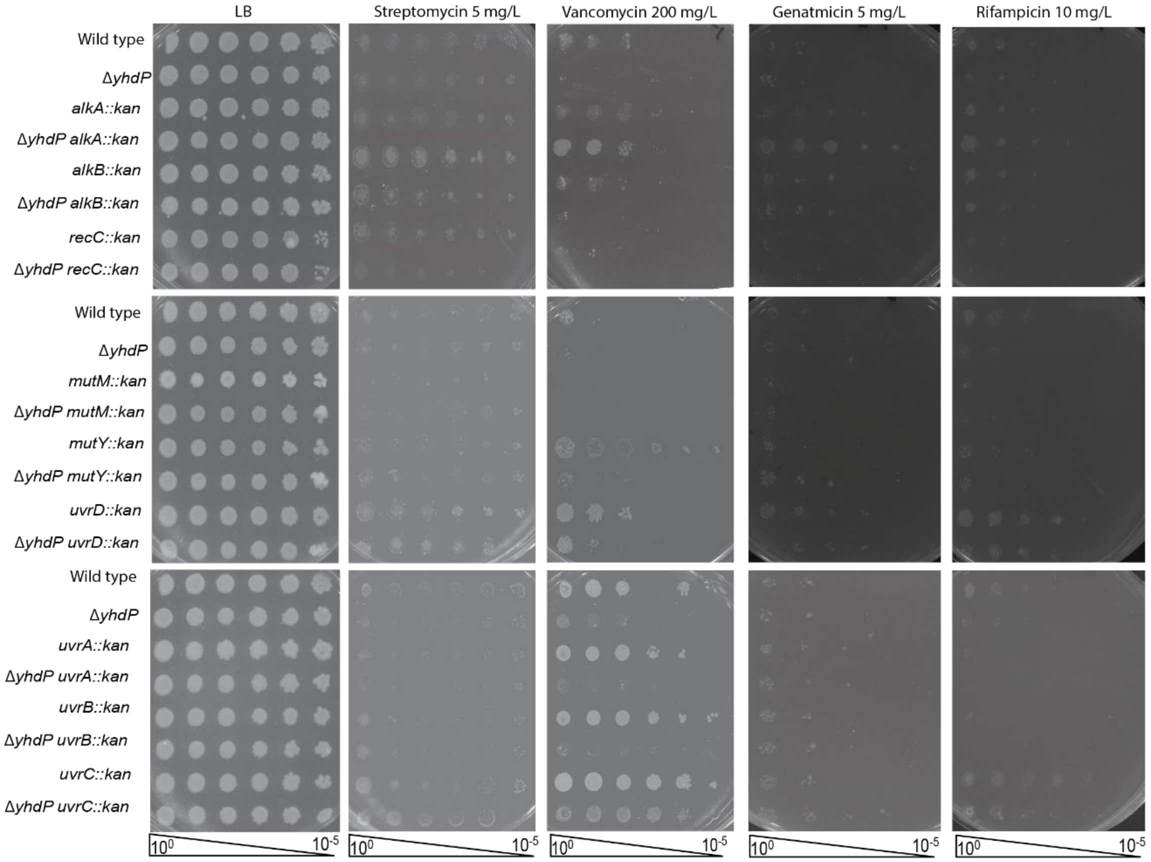


**Fig. S3. Increased antibiotic resistance does not occur with the inhibition of other DNA repair pathways.** To investigate whether the increased antibiotic resistance we observe with loss of MMR occurs with other DNA repair pathways, we made deletions of several DNA repair associated genes that span multiple DNA repair mechanisms (nucleotide excision, base excision, aklyation, etc.) and we found inhibition of other DNA repair pathways does not lead to a similar pattern of increased resistance to that observed with loss of MMR.

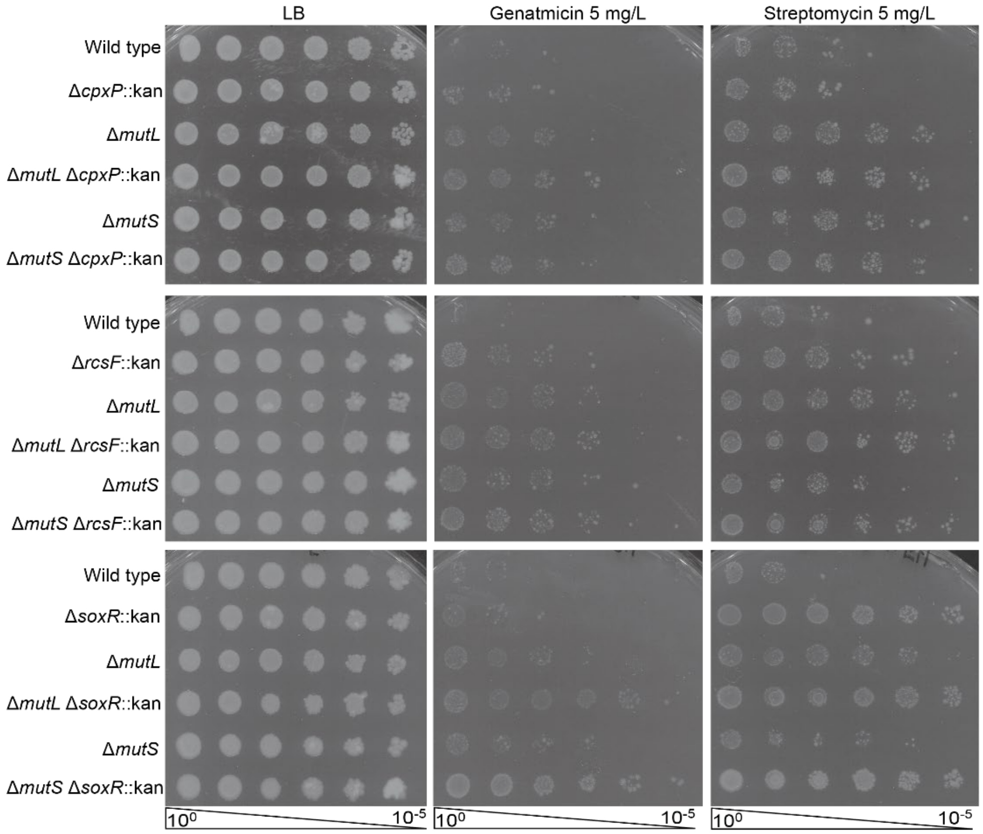


**Fig. S4. Increased antibiotic resistance with loss of MMR is independent of several stress responses.** Deletions in various known stress response pathways were made to see whether they played a role in increased resistance in Δ*mutL* or Δ*mutS* strains. If these stress responses were involved in the increased resistance we observe with loss of MMR, we would expect decreased resistance when the stress response is inhibited (*rcsF*::kan, *soxR*::kan) or increase resistance with its activation (*cpxP*::kan). No such changes were observed.

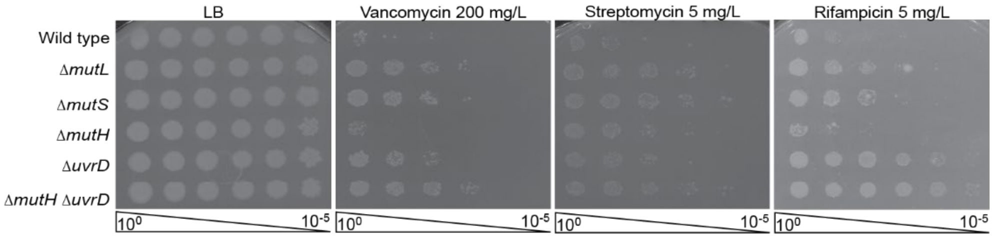


**Fig. S5. Increased resistance from loss of MMR correlates with reported changes in homologous recombination rates.** Resistance from loss of MutL and MutS is greater than that of MutH or UvrD. However, resistance from loss of both MutH and UvrD is similar to that of MutL and MutS. These changes correlate well with the reported effects of MMR mutants on rates of homologous recombination.

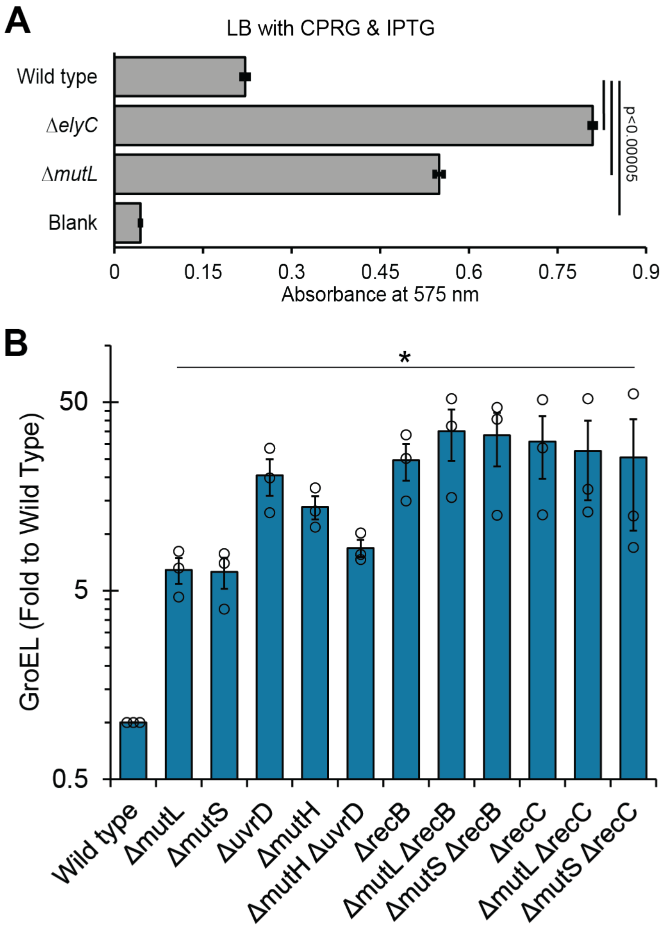


**Fig. S6. Loss of MMR causes increased lysis. (A)** The indicated strains were grown overnight with IPTG and then culture supernatants were harvested and incubated with CPRG before chlorophenol red absorbance was assayed to quantitate CPRG activity. An Δ*elyC* strain serves as a positive control. The Δ*mutL* strain showed increased CPRG activity compared to the wild type. Data is representative of at least 3 independent experiments. Mean ± Std Dev is shown. **(B)** Quantification of GroEL levels in the culture supernatant of the indicated strains. Data are the mean of three biological replicates ± the SEM. * $p<0.05$ by Mann Whitney test compared to wild type.

**Table S1. Loss of MMR increases antibiotic minimum inhibitory concentration**

| Strain | Minimum Inhibitory Contraction (mg/L)[a] | |
|---|---|---|
| | Gentamicin | Streptomycin |
| Wild Type | 3.6 | 3.9 |
| Δ*mutL* | 9.4[b] | 9.8[b] |
| Δ*mutS* | 8.2[b] | 12.4[b] |

[a] Values are shown as geometric means of three biological replicates.

[b] $p<0.05$ vs. wild type by Mann-Whitney test

**Table S2. Strains used in this study**

| Strain | Genotype | Reference |
|---|---|---|
| MG1655 | K-12 F$^-$ λ$^-$ *rph-1* | (1) |
| DB033 | MG1655 Δ*mutL* | This study |
| DB035 | MG1655 Δ*mutS* | This study |
| DB038 | MG1655 Δ*acrA* Δ*mutL* | This study |
| DB039 | MG1655 Δ*acrA* Δ*mutS* | This study |
| DB041 | MG1655 Δ*pldA* | This study |
| DB043 | MG1655 Δ*yhdP* Δ*mutL* | This study |
| DB044 | MG1655 Δ*yhdP* Δ*mutS* | This study |
| DB046 | MG1655 Δ*mlaA* | This study |
| DB063 | MG1655 Δ*mlaA* Δ*mutL* | This study |
| DB064 | MG1655 Δ*mlaA* ΔmutS | This study |
| DB069 | MG1655 Δ*pldA* Δ*mutL* | This study |
| DB070 | MG1655 Δ*pldA* Δ*mutS* | This study |
| DB081 | MG1655 Δ*mutH* | This study |
| DB691 | MG1655 *lexA3*(Ind-) *malF3089*::Tn*10* | This study |
| DB692 | MG1655 Δ*bamE* *lexA3*(Ind-) *malF3089*::Tn*10* | This study |
| DB693 | MG1655 Δ*bamE* Δ*mutL* *lexA3*(Ind-) *malF3089*::Tn*10* | This study |
| DB121 | MG1655 Δ*bamB*:kan | This study |
| DB122 | MG1655 Δ*mutL* *bamA101*::Tn*5* | This study |
| DB124 | MG1655 Δ*mutS* *bamA101*::Tn*5* | This study |
| DB162 | MG1655 *bamA101*::Tn*5* | This study |
| DB163 | MG1655 Δ*bamB*::kan | This study |
| DB164 | MG1655 Δ*bamB* Δ*mutL* | This study |
| DB165 | MG1655 Δ*bamB* Δ*mutS* | This study |
| DB445 | MG1655 Δ*clpP*::kan | This study |
| DB446 | MG1655 Δ*mutL* Δ*clpP*::kan | This study |
| DB447 | MG1655 Δ*mutS* Δ*clpP*::kan | This study |
| DB454 | MG1655 Δ*soxR*::kan | This study |
| DB455 | MG1655 Δ*mutL* Δ*soxR*::kan | This study |
| DB456 | MG1655 Δ*mutS* Δ*soxR*::kan | This study |
| DB467 | MG1655 Δ*rcsF::*kan | This study |
| DB468 | MG1655 Δ*mutL* Δ*rcsF*::kan | This study |
| DB469 | MG1655 Δ*mutS* Δ*rcsF*::kan | This study |
| DB507 | MG1655 Δ*recA*::cm | This study |
| DB508 | MG1655 Δ*yhdP* Δ*recA*::cm | This study |
| DB509 | MG1655 Δ*bamE* Δ*recA*::cm | This study |
| DB510 | MG1655 Δ*mutL* Δ*recA*::cm | This study |

| | | |
|---|---|---|
| DB511 | MG1655 Δ*yhdP* Δ*mutL* Δ*recA*::cm | This study |
| DB512 | MG1655 Δ*bamE* Δ*mutL* Δ*recA*::cm | This study |
| LT2 | *Salmonella enterica* subsp. *enterica* serovar Typhimurium strain LT2 | ATCC 700720 |
| DB619 | LT2 Δ*mutL::*kan | This study |
| DB620 | LT2 Δ*mutS*::kan | This study |
| DB602 | MG1655 Δ*elyC* | This study |
| DB649 | MG1655 Δ*recB*::kan | This study |
| DB651 | MG1655 Δ*mutL* Δ*recB*::kan | This study |
| DB652 | MG1655 Δ*mutS* Δ*recB*::kan | This study |
| DB667 | MG1655 Δ*recC*::Kan | This study |
| DB668 | MG1655 *△mutL* Δ*recC*::kan | This study |
| DB669 | MG1655 *△mutS* Δ*recC*::kan | This study |
| AM043 | MG1655 Δ*acrA*::kan | This study |
| AM044 | MG1655 Δ*acrB*::kan | This study |
| AM102 | MG1655 Δ*acrA* | (2) |
| AM103 | MG1655 Δ*acrB* | (2) |
| AM452 | MG1655 Δ*yhdP* Δ*mutL* | This study |
| AM453 | MG1655 Δ*yhdP* Δ*mutS* | This study |
| AM182 | MG1655 Δ*yhdP* | (2) |
| AM449 | MG1655 Δ*bamE* | This study |
| DB029 | MG1655 Δ*mutL* Δ*bamE* | This study |
| DB031 | MG1655 Δ*mutS* Δ*bamE* | This study |
| DB561 | MG1655 Δ*dam*::kan | This study |
| DB126 | MG1655 pBAD33 | This study |
| DB128 | MG1655 pBAD33-*mutL* | This study |
| DB130 | MG1655 pBAD33-*mutS* | This study |
| DB138 | MG1655 Δ*mutL* pBAD33 | This study |
| DB140 | MG1655 Δ*mutL* pBAD33-*mutL* | This study |
| DB142 | MG1655 Δ*mutS* pBAD33 | This study |
| DB144 | MG1655 Δ*mutS* pBAD33-*mutS* | This study |

**Table S3. Primers used in this study**

| Name | Primer |
|---|---|
| mutL Fwd homology | CCAGTACCAACCAGCCTGGCGCCATAACCGCCGCGAATTAAG GAGATTTCATTCCGGGGATCCGTCGACC |
| mutL Rev homology | TCGCCTTAGGCAGGCTCGCCTTGCTTACAtcaTTCATGCTTCAGG GCGTTGTGTAGGCTGGAGCTGCTTCG |
| mutS Fwd homology | CCTGGGGCCACATGAGCACTTGAAAGAATAAAAAAACGAGTAAA ATCAATATTCCGGGGATCCGTCGACC |
| mutS Rev homology | ATAGCGCGGCGCTATCGGGTCTGACACGTttaCACCAGACTTTTC AGCCGGTGTAGGCTGGAGCTGCTTCG |
| Del-recA F | CAGAACATATTGACTATCCGGTATTACCCGGCATGACAGGAGTA AAAATGTGTAGGCTGGAGCTGCTTCG |
| Del-recA cm R | ATGCGACCCTTGTGTATCAAACAAGACGATTAAAAATCTTCGTTAG TTTCCATATGAATATCCTCCTTAG |
| pBAD_fwd | CCGGGGATCCTCTAGAGTC |
| pBAD_rev | GTACCGAGCTCGAATTCG |
| mutL_fwd | AGCGAATTCGAGCTCGGTACGATCGCACGCTGCCAAAC |
| mutL_rev | CGACTCTAGAGGATCCCCGGTCACTCATCTTTCAGGGCTTTTATC |
| mutS_fwd | AGCGAATTCGAGCTCGGTACATTTAATATCAGGGAACCGG |
| mutS-rev | CGACTCTAGAGGATCCCCGGTTACACCAGGCTCTTCAAG |
